# Seq-InSite: sequence supersedes structure for protein interaction site prediction

**DOI:** 10.1101/2023.06.19.545575

**Authors:** SeyedMohsen Hosseini, G. Brian Golding, Lucian Ilie

## Abstract

Proteins accomplish cellular functions by interacting with each other, which makes the prediction of interaction sites a fundamental problem. Computational prediction of the interaction sites has been studied extensively, with the structure-based programs being the most accurate, while the sequence-based ones being much more widely applicable, as the sequences available outnumber the structures by two orders of magnitude. We provide here the first solution that achieves both goals. Our new sequence-based program, Seq-InSite, greatly surpasses the performance of sequence-based models, matching the quality of state-of-the-art structure-based predictors, thus effectively superseding the need for models requiring structure. Seq-InSite is illustrated using an analysis of four protein sequences. Seq-InSite is freely available as a web server at `seq-insite.csd.uwo.ca` and as free source code, including trained models and all datasets used for training and testing, at github.com/lucian-ilie/seq-insite.

## 1 Introduction

Proteins are among the most influential molecules in a cell, being responsible for many functions including cell growth, gene expression, and intercellular communication [54]. Proteins interact with each other for proper function, therefore it is crucial to study them in the context of their interacting partners. These interactions involve the binding of two or more proteins to form a complex structure and detecting them will ultimately help researchers understand the mechanism of various biological processes, disease development, and drug design [23, 54].

Observing how proteins interact may lead to discovery of previously unknown proteins. Additionally it could also reveals an unforeseen functionality of a protein [35]. In order to fully comprehend the molecular mechanisms of protein-protein interactions, it is essential to identify the specific residues within proteins that facilitate the formation of interactions. Protein interactions can be detected experimentally or predicted by computational methods. Although experimental methods like immunoprecipitation [26], pull-down assay [27], surface plasmon resonance [12], bacterial two-hybrid [22] and cytology two-hybrid [3] may generate a more accurate description of interaction sites, they are expensive and time consuming. Computational methods are fast and cheep and can provide useful information to be used as such or to generate the most likely candidates to be confirmed experimentally. Improving the quality of computational predictions is thus a fundamental problem.

Protein interaction site prediction is the problem of predicting the locations of a given protein where interactions are likely to take place. It involves using bioinformatics algorithms and machine learning techniques to analyze large amounts of protein interaction data. Various machine learning techniques have been employed for this problem, such as logistic regression [55], random forest [6], and neural networks [18], etc. In spite of considerable progress, much work is needed to achieve the necessary accuracy that can have a great impact on the understanding of the molecular mechanisms.

Computational models can be classified into two large categories, according to the type of information used as input: sequence or structure. Structured-based methods include the use of molecular dynamics simulations that provides insight into the dynamics of protein interactions [8] or the three-dimensional structures of proteins to predict interaction sites [52, 49, 53]. The main drawback of these methods is the limited availability of protein’s structure. On the other hand, sequence-based methods [18, 25, 55] rely solely on the protein sequences, thus having the advantage of being applicable to a far broader range of proteins, as the number of available protein sequences outnumbers that of structures by two orders of magnitude [38].

The majority of models utilize supervised feature extractions in order to create some form of representation for each amino acid including position-specific scoring matrix (PSSM), hidden Markov models (HMM) [41], and a dictionary of protein’s secondary structure (DSSP) [20].

Among the most successful structure-based models, DeepP-PISP [53] combines sequential information with structural information generated by DSSP to predict interacting sites. Its architecture is divided into two sections, global and local.

The global part employs a fully connected layer to embed the one-hot-encoding representation of amino acids and the embedding output concatenated to the other features and fed to a TextCNN. The local part uses a sliding window with an odd size centered on the site considered for prediction. Another successful structure-based program, GraphPPIS [52], employs a deep graph convolutional network. The graph of each sequence, created using the PDB [5] file of the protein, and the computed features are fed to a graph convolutional network with residual connections. The RGN program [49] attempts to improve the performance of GraphPPIS by adding a Bert language model [10].

From the top sequence-based models, SCRIBER [55] uses logistic regression, DLPred[54] uses an SLSTM, which is a modified LSTM, DELPHI [25] employs an ensemble model that incorporates RNN and CNN with many features, including a novel embedding based on ProtVec [2], and PITHIA [18] using the embeddings computed by the MSA-transformer [40] with an attention-based architecture.

The rise of embeddings in natural language processing, from contextual independent word2vec [31] or GloVe[36] to the more effective context dependent models such as ELMo [37] and Bert [10] had implications in the field of proteomics as well, where protein embeddings have been produced using the above mentioned models [4, 33, 14, 40].

In our previous models, we employed a simplified ProtVec in DELPHI, in addition to other features, while in PITHIA, the use of the MSA (multiple sequence alignment) transformer embeddings obviated the need for the traditional features. In spite of great performance, one of the main problems of PITHIA, AlphaFold [19], the MSA-transformer, or any other model that utilizes MSA information, is that high quality multiple sequence alignment are of paramount importance for the proper performance of the model. Additionally, these models require a good amount of resource and time in order to create the MSA for each given protein. ProtT5-XL is a model presented in Prottrans [14] in an attempt to compute embeddings without using MSA information. ProtT5-XL employs a T5 architecture with some changes including the same masking procedure that was employed by the Bert architecture to calculate a representation for each amino acid in the sequence. The ProtT5-XL training procedure is to train a T5 model on the BFD dataset [46, 44] and then fine tune it on Uniref50 [47].

In this paper we introduce our new model, Seq-InSite (Sequence-based Interaction Site), that uses both embeddings, computed by ProtT5-XL and the MSA-transformer, in an ensemble architecture model that combines a multi-layer perceptron (MLP) and an LSTM. The result is a model that surpasses the current state-of-the-art sequence-based programs by a wide margin and performs on par with or better than the current state-of-the-art structure-based models. This is proved by thorough testing and comparison with top models on the most widely used datasets. In addition, Seq-InSite is applied to four protein sequences. Seq-InSite makes it possible to obtain structure level quality predictions for the widely available protein sequences. In order to make it as widely available as possible, we provide access to the program as a web server as well as as free source code, including trained models and datasets.

Concurrently with our development, the group of Martelli produced a very competitive program, ISPRED-SEQ [30]. Similarly with PITHIA, their program uses embeddings only as input features, namely Prot-T5 and ESM1-b [42].

## 2 Materials and Methods

### 2.1 Datasets

We use three benchmark datasets have been used frequently in the field, Dset_186, _72 [32], and Dset_164 [11], as well as a recent dataset, Dset_448, created by Zhang and Kurgan [55]; Dset_355 is a subset of Dset_448 which is described in the Results section.

In contrast to sequence-based models such as SCRIBBER, DLPred, DELPHI, and PITHIA which have their own training dataset, structure-based models like DeepPPISP, GraphPPIS, and RGN combine the above three datasets and divide their union into training and testing. For example, DeepP-PISP uses a test set of 70 proteins while GraphPPIS created a different subset of 60 proteins as its test set; we are using these datasets as well for testing, and they are called Dset_60 and Dset_70.

GraphPPIS introduced also a different dataset, Test 315, created based on the newly solved protein complexes in PDB (January 2014 – May 2021) by removing any overlapping or similar protein structures with any proteins in the Dset_ 186, Dset_ 72, or Dset_ 64 datasets. For uniform naming, we shall call Test_315 as Dset_ 315.

#### 2.1.1 Training and validation datasets

The training dataset was produced as follows. We took all 22,654 proteins from the most recent version of PiSite [17] (January 2019) and removed all sequences with no interaction residues or containing less than 50 amino acids; 14,203 sequences remained after this elimination. We then used PSI-CD-HIT [24, 15] to remove any sequences that have any similarity above 25%, as customary in this area, with any of the widely used testing datasets, Dset_ 72, Dset_ 164, Dset_ 186, and Dset_ 448. The remaining proteins, 11,523, form Seq-InSite’s training dataset; this is further split into training (99%) and validation (1%) sets; this size of validation data is sufficient given that our dataset has about 2.9 million samples.

Table 1 gives an overview of all these datasets.

**Table 1:** Datasets overview; all but the last dataset are used for testing.

| Dataset | Proteins<br>sequences | Mean<br>length | Total<br>residues | Interacting<br>residues | % |
| --- | --- | --- | --- | --- | --- |
| Dset_448 | 448 | 260 | 116,500 | 15,810 | 13.60 |
| Dset_355 | 355 | 270 | 95,940 | 11,467 | 11.95 |
| Dset_315 | 315 | 208 | 65,331 | 9,355 | 14.32 |
| Dset_186 | 186 | 210 | 39,008 | 6,128 | 15.70 |
| Dset_164 | 164 | 223 | 36,589 | 5,276 | 14.40 |
| Dset_72 | 72 | 211 | 15,236 | 3,832 | 25.20 |
| Dset_70 | 70 | 168 | 11,791 | 2,332 | 19.78 |
| Dset_60 | 60 | 219 | 13,144 | 2,075 | 15.78 |
| Train+Validate | 11,523 | 252 | 2,902,667 | 545,724 | 18.80 |

### 2.2 Input embeddings

The transformer architecture has been successful in producing superior protein folding models, including the highly appraised AlphaFold [19], which employs multiple sequence alignment in a supervised manner. MSA-transformer [40] also uses attention and multiple sequence alignment to generate powerful 768-dimensional amino acid embeddings in an unsupervised manner. The T5 language model [39], a variant of the transformer architecture, has demonstrated its effectiveness in large-scale natural language processing and has shown that the simplification of models like BERT can result in lower performance. ProtT5-XL [14] uses a T5 architecture with the Bert masking procedure to compute amino acid embeddings. It trains a T5 model with 32 heads on the BFD dataset [46, 44] for 1.2M epochs and then fine tunes it on Uniref50 [47] for 991k more epochs and AdaFactor optimizer with inverse square root learning rate schedule were utilized for training. ProtT5-XL embeddings have size 1024.

ProtT5-XL is particularly useful for proteins with long sequences or those that do not have a high-quality multiple sequence alignments, thanks to its gigantic and state-of-the-art architecture that can represent any amino acid with a high-quality embedding. On the other hand, MSA-transformers can capture more complex relationships between protein sequences that have a high-quality alignments by utilizing the existing relations among the aligned protein sequences. Prot-trans also developed a larger model, but increasing the size did not improve its performance. Our tests showed that ProtT5-XL is the superior model for predicting interaction sites.

Our new program, Seq-InSite, uses both MSA-transformer and ProtT5-XL embeddings as inputs to create a more comprehensive and powerful model for predicting protein interaction sites. The combination of the two architectures allows Seq-InSite to better capture the complexity of protein sequences and achieve better performance. The architecture of Seq-InSite is described in detail in the next section.

### 2.3 Model architecture

Seq-InSite’s architecture is inspired by ensemble learning, with two MLP and LSTM components that represent an ensemble of both ProtT5-XL and MncatSA-transformer embeddings. These components are fused together through multiple dense layers to predict protein interaction sites. The architecture is depicted in Figure 1 and the parameters are given in Table 2.

**Table 2:** Parameters of the model.

| Model Parameters |  | Values |
| --- | --- | --- |
| MLP | Layer sizes | $1024 \times 9, 768 \times 9, 256, 128, 16, 1$ |
| RNN | LSTM unites | 64 |
|  | Fully connected unit | 64, 1 |
|  | Activation function | tanh |
| All | Batch size | 1024 |
|  | Dropout rate | 0.3 |
| | Optimizer | Adam ($\beta_1 = .9; \beta_2 = .999$) |
|  | Loss function | Binary cross entropy |
|  | Learning rate | 0.001 |
|  | Patience (early stop) | 4 |

**Figure 1:**
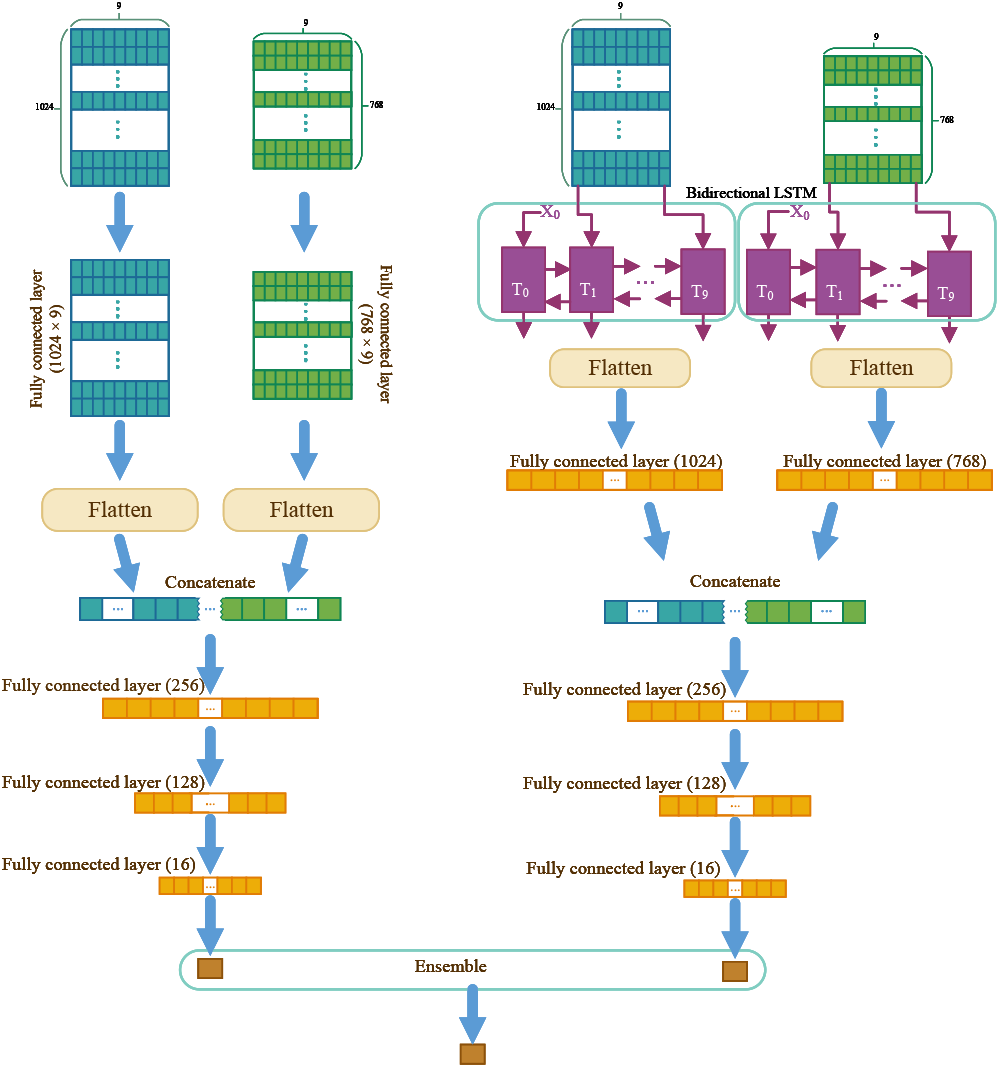
The architecture of Seq-InSite.

Seq-InSite uses a many-to-one structure, where a window centred on each target aminoer acid is used to gather information from neighbouring residues in an attempt to predict interaction propensity. The window size, *w* = 9, has been determined experimentally. The sequence’s ends are padded with zeros.

#### 2.3.1 Architecture of MLP Network

The MLP component, shown in the left side of Figure 1, contains a fully connected layer, a flatten layer, and four fully connected layers with dropout. The input is of size *w×e*, where *w* is the window size and *e* is the embedding size that depends on the language model variant being used (MSA-transformer = 768 or ProtT5-XL = 1024), and employs the sigmoid activation function in the final layer to generate output probabilities, while all other layers use the ReLU activation function.

#### 2.3.2 Architecture of RNN Network

The RNN component of Seq-InSite, depicted on the right side of Figure 1, employs a bidirectional LSTM layer with 64 units, followed by a flatten layer and two fully connected layers that reduce the dimensionality to one, with input size varying depending on the language model variant being used, and the LSTM layer set to return a sequence of values.

#### 2.3.3 Architecture of Ensemble Network

The final model employs an ensemble network to combine information from both the MLP and LSTM components to predict the interaction propensity of each amino acid in a protein sequence. The MLP and LSTM branches process the input data separately, using MSA-transformer and ProtT5-XL, and produce their own predictions. The final prediction is obtained by averaging the predictions from the two branches to improve accuracy and reliability. The architecture of the model is shown in Figure 1.

### 2.4 Implementation

The code is implemented using Keras [7] with TensorFlow GPU [1] as the backend. It utilizes generators to handle the large data, for which reason local shuffling is used, since generators read small parts of the fasta file at a time. A batch size of 1024 is used to enable multiple proteins to be included in each batch. By using generators, we significantly reduced the memory requirement. The program requires 110GB of RAM, and the training process takes about 25 minutes per epoch. During testing, the program takes about 18 seconds to process a sequence if embeddings are already available. Computing MSA-transformer embeddings takes 10-20 minutes; ProtT5-XL embeddings require approximately 40 seconds.

### 2.5 Model selection

Multiple models, involving combinations of MLP, RNN (LSTM), CNN, and Transformer architectures, were constructed and evaluated using the training dataset. No testing data was used to choose the final model. Each model was trained on 99% of the available data, and its performance was evaluated using the remaining 1% validation data; see Table 1. The final Seq-InSite model was selected based on its performance, as measured by the area under the PR curve, on the validation data. Since our data is skewed, the PR curve represents better the performance [9], and the area under the curve indicates overall performance, which is not threshold dependent.

Notably, no protein sequences used for model selection come from the primary datasets, namely Dset_ 72, Dset_ 164, Dset_ 186, and Dset_ 448. In fact, not a single protein sequence that shares more than 25% similarity with any of the proteins in these datasets was used for selecting the final model.

## 3 Results

### 3.1 Competing methods

Despite relying solely on sequence data to predict interaction sites, we compared our model not only to sequence-based methods but also to state-of-the-art structure-based methods. We selected for comparison a number of methods based on the code availability, good performance reported in their papers, or having received significant attention in the literature. These include sequence-based models: CRFPPI [51], DELPHI [25], DLPred [54], ISPRED-SEQ [30], LORIS [11], PITHIA [18], PSIVER [32], SCRIBER [55], SPRINGS [43], SPRINT [48], and SSWRF [50]; and structure-based models: AttentionCNN [28], DeepPPISP [53], EGRET [29], GraphP-PIS [52], HN-PPISP [21], MaSIF [16], and RGN [49].

However, problems encountered while attempting to run the code of some of the methods above prevented us from performing the full comparison on all datasets. Ideally, we would have liked to train and test all methods ourselves, for the most reliable comparison. Instead, in some instances, we had to reply upon pre-trained code or the figures provided in the corresponding papers, which is less than optimal.

We could not run AttentionCNN [28], EGRET [29], and HN-PPISP [21], and our requests to the corresponding authors were not answered.

For RGN [49], the (pre-trained) code provided in the website performed less well than claimed in their paper. The authors responded to our request and provided a different pretrained model, which we ran and reported its results, which are still not as good as reported in their paper. Our request for a source code which we could train ourselves was not answered.

We could not properly include the results from the MaSIF [16] program due to significant similarities between its training data and the testing ones.

More details are given below for each dataset involved.

### 3.2 Evaluation metrics

It is good practice in binary classification evaluation to use many metrics. We employ sensitivity (or recall), precision, specificity, accuracy, F1-score, Matthew’s correlation coefficient (MCC), area under the receiver operating characteristic curve (ROC), and area under the precision-recall curve (PR). We briefly recall their definitions, where *TP* represents the number of correctly predicted interaction sites, *TN* represents the number of correctly predicted non-interaction sites, *FP* represents the number of incorrectly predicted interaction sites, and *FN* represents the number of incorrectly predicted non-interaction sites:

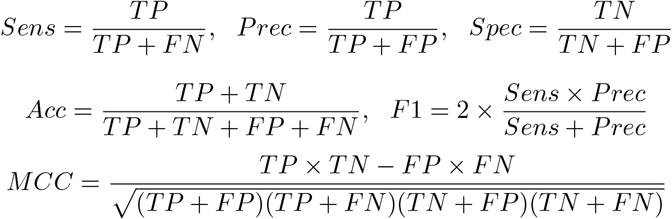

Except for the areas under the ROC and PR curves, all the aforementioned metrics are dependent on the threshold in order to distinguish positive and negative predictions. According to common practice used in many previous studies, the threshold was set such that the number of predicted interaction residues equals the number of actual interaction residues. As a result, *TP* + *FP* = *TP* + *FN*, which implies that *FP* = *FN*, resulting in the equality of sensitivity, precision, and F1-score.

### 3.3 Performance Comparison

Most sequence-based methods for predicting protein interactions are developed with their own training dataset, with Dset_ 186, Dset_ 164, Dset_ 72, and Dset_ 448 being common test sets used to compare the performance of different methods. However, as the number of available protein structures is limited, structure-based methods face the challenge of finding a good quality dataset for training procedure. This limitation resulted in using the 422 proteins in datasets Dset_ 186, Dset_ 164, and Dset_ 72 to both train and test. For instance, DeepPPISP, EGRET, and HN-PPISP use a test set consisting of 70 out of the 422 available proteins and the 352 remaining protein as training. Similarly, GrapPPIS and RGN make use of a test set of 60 proteins, training on the rest. Due to the availability of the methods we had to focus on Dset_ 60 as a representative to the three traditional dataset. We consider Dset_ 70 also, but in a more limited way.

In addition to the above datasets, we consider also Dset_ 315, introduced by GraphPPIS, as further comparison with structure-based methods.

All comparisons we perform have as their goal not only to indicate the best performing method in a given test, but rather to compare the differences in performance. To that end, we present all tables as heat maps, for better visual interpretation.

#### 3.3.1 Comparison against sequence-based models

Dset_ 448 is among the most recent datasets, that has gained attention in the field. Because the training set of DLPred, one of the main sequence-based competitors, shares significant similarities with Dset_ 448, a subset of Dset_ 448 was created by Delphi [25], called Dset_ 335, which contains all 355 proteins from Dset_ 448 that have no significant similarity with the training set of DLPred. Dset_ 355, which we already include in Table 1, is used in place of Dset_ 448 in our comparisons.

We use Dset_ 355 to compare Seq-InSite with several sequence-based models. The comparison is presented in Table 3, with ROC and PR curves plotted in Figure 2(a)-(b). Both the table and the curves indicate very clearly that the performance of the Seq-InSite model is far better than all the other models. Specifically, when evaluating the area under the precision-recall curve, Seq-InSite shows a remarkable improvement of 52% over the second best model. Additionally, Seq-InSite demonstrates a 53% improvement on Matthews correlation coefficient (MCC) compared to the second best model.

**Table 3:** Performance comparison on Dset_ 355. All methods included in the table are sequence-based. Darker colour indicates better performance. The corresponding ROC and PR curves are shown in Figure 2(a)-(b).

| Model | Sens | Spec | Prec | Acc | F1 | MCC | ROC | PR |
| --- | --- | --- | --- | --- | --- | --- | --- | --- |
| CRFPPI | 0.245 | 0.898 | 0.245 | 0.820 | 0.245 | 0.143 | 0.662 | 0.214 |
| DELPHI | 0.368 | 0.914 | 0.368 | 0.849 | 0.368 | 0.282 | 0.749 | 0.333 |
| DLPred | 0.305 | 0.906 | 0.305 | 0.834 | 0.305 | 0.211 | 0.725 | 0.268 |
| LORIS | 0.240 | 0.897 | 0.240 | 0.818 | 0.240 | 0.137 | 0.637 | 0.203 |
| PITHIA | 0.384 | 0.916 | 0.384 | 0.853 | 0.384 | 0.301 | 0.769 | 0.350 |
| PSIVER | 0.177 | 0.888 | 0.177 | 0.803 | 0.177 | 0.065 | 0.583 | 0.155 |
| SCRIBER | 0.322 | 0.908 | 0.322 | 0.838 | 0.322 | 0.230 | 0.719 | 0.275 |
| SPRINGS | 0.209 | 0.893 | 0.209 | 0.811 | 0.209 | 0.102 | 0.608 | 0.178 |
| SPRINT | 0.167 | 0.887 | 0.167 | 0.801 | 0.167 | 0.054 | 0.571 | 0.150 |
| SSWRF | 0.267 | 0.901 | 0.267 | 0.825 | 0.267 | 0.168 | 0.667 | 0.228 |
| Seq-InSite | 0.525 | 0.935 | 0.525 | 0.886 | 0.525 | 0.460 | 0.860 | 0.533 |

**Figure 2:**
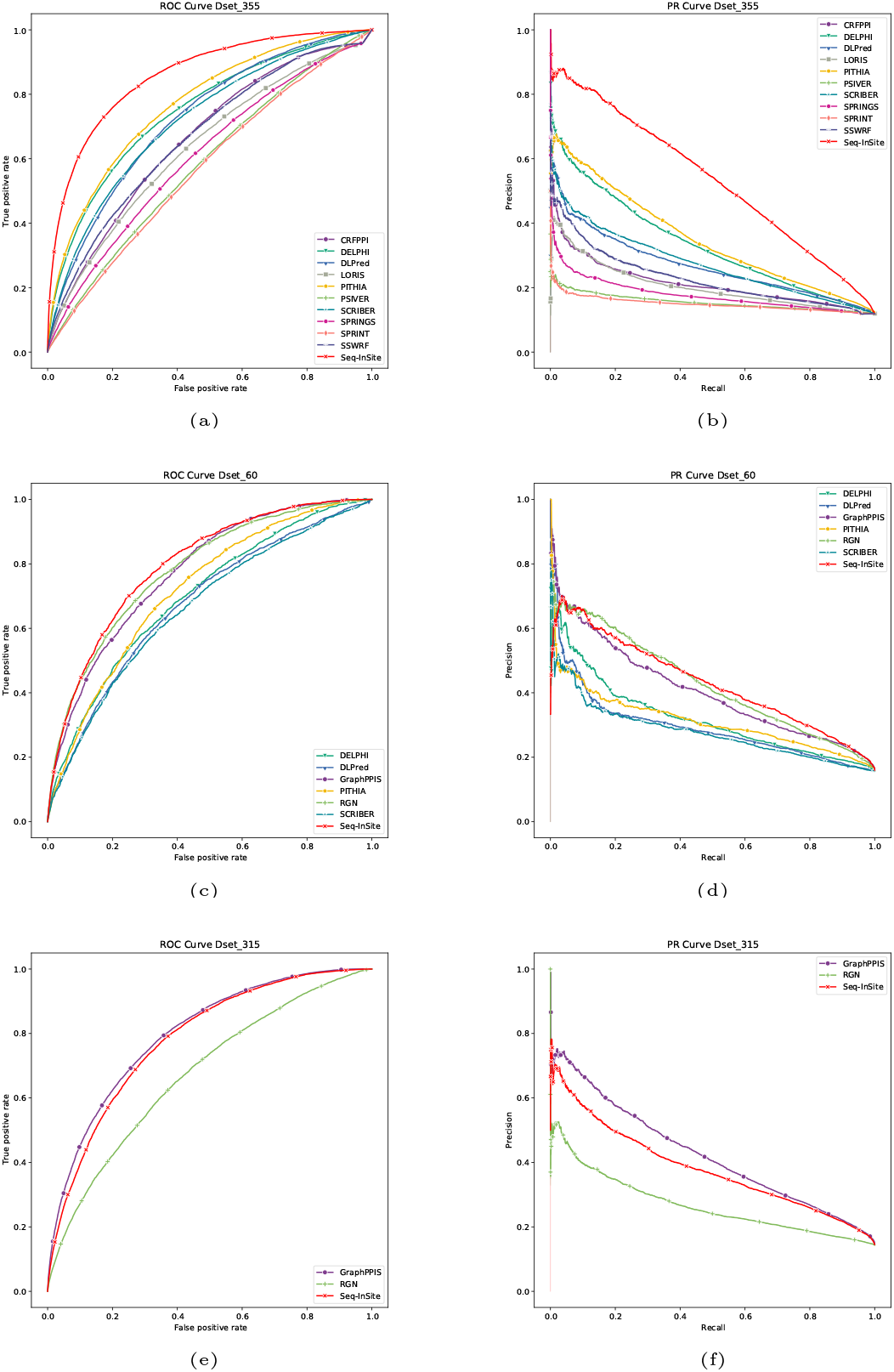
ROC and PR curves for the tests sets Dset_ 355, Dset_ 60, and Dset_ 315. The area under the curves is given in Tables 3, 5, and 6, resp. The left column contains the ROC curves and the right one the PR curves. The two curves for the same dataset are in the same row.

Regarding the very recent ISPRED-SEQ program, we can perform only a partial comparison, since its source code is not available. Using the numbers reported in the paper [30], on dataset Dset_ 355, ISPRED-SEQ has F1-score 0.46, MCC 0.39, and area under the ROC curve 0.82. Seq-InSite exceeds these values by 14%, 20%, and 5%, resp. Unfortunately, our most important metric, the area under the PR-curve, has not been provided. Also, without the source code, we cannot add its ROC and PR curves to Figure 2. In spite of this, ISPRED-SEQ appears to perform second best, after Seq-InSite, among all sequence-based programs. This is supported as well by comparison on Dset_ 448, which we used for the ablation study below, which indicates similar behaviour with Dset_ 355.

#### 3.3.2 Comparison against structure-based models

Due to the limitations explained earlier, we need to find a way to properly train our model for testing on datasets Dset_ 60, Dset_ 70, and Dset_ 315, such that we include as many protein sequences as possible in training, without having any similarity (above 25%) with testing data. Therefore, we had to adjust our trainign dataset for each of the three testing datasets. In case case, we started with the 14,203 proteins computed earlier (see Training and validation datasets) and removed all protein sequences that have more than 25% similarity with any sequence in the testing datasets. The obtained training datasets are shown in Table 4. Each set is split into 99%/1% training and validation.

**Table 4:**
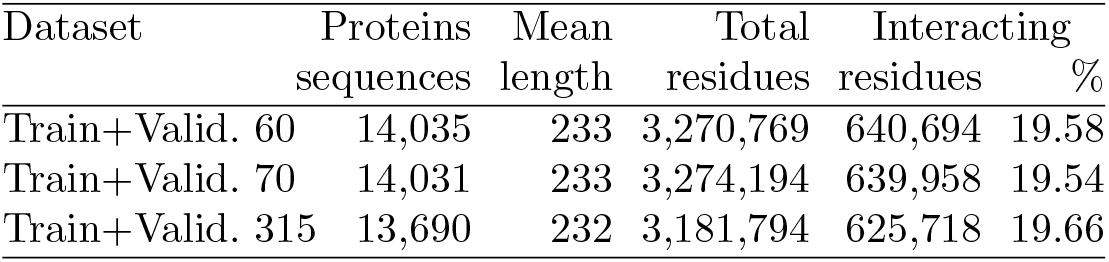
Datasets used for training in order to test on Dset_ 60, Dset_ 70, and Dset_ 315.

**Table 5:**
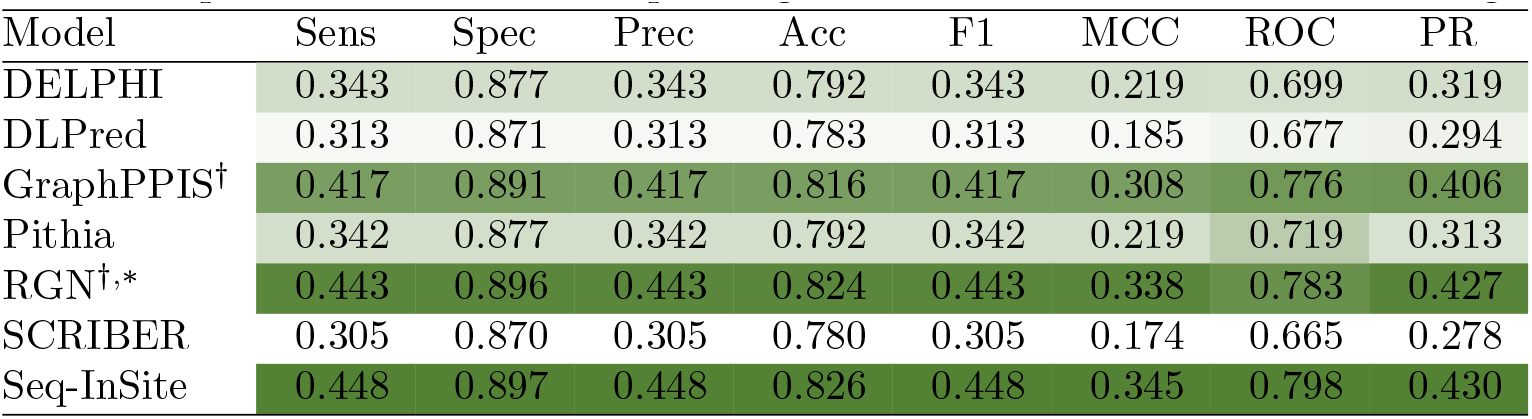
Performance comparison on Dset_ 60. The methods marked by ^†^ are structure-based; the rest are sequence-based. The methods marked by ^*^ were not trained by us, pre-trained code was used; the remaining methods were trained by us. Darker colour indicates better performance. The corresponding ROC and PR curves are shown in Figure 2(c)-(d).

**Table 6:** Performance comparison on Dset_ 315. The methods marked by ^†^ are structure-based; the rest are sequence-based. The methods marked by ^*^ were not trained by us, pre-trained code was used; the remaining methods were trained by us. Darker colour indicates better performance. The corresponding ROC and PR curves are shown in Figure 2(e)-(f).

| Model | Sens | Spec | Prec | Acc | F1 | MCC | ROC | PR |
| --- | --- | --- | --- | --- | --- | --- | --- | --- |
| GraphPPIS† | 0.437 | 0.906 | 0.437 | 0.839 | 0.437 | 0.343 | 0.798 | 0.423 |
| RGN$\dagger,*$ | 0.346 | 0.891 | 0.346 | 0.813 | 0.346 | 0.237 | 0.711 | 0.316 |
| Seq-InSite | 0.398 | 0.899 | 0.398 | 0.827 | 0.398 | 0.297 | 0.782 | 0.380 |

##### Comparison on Dset_ 60

The comparison on Dset_ 60 is presented in Table 5, with ROC and PR curves plotted in Figure 2(c)-(d). This comparison is very interesting because we managed to include both sequence-based and structure-based methods. The most striking feature, evident from both the dark rows in Table 5 and the curve grouping in Figure 2(c)-(d), is the difference between sequence-based models and structure-based models, with the latter having a clear advantage. Our new method, Seq-InSite, is the exception, clearly belonging to the higher group, as the top performing method. Seq-InSite has the highest area under both the PR curve and the ROC curve. It is important to mention that for the RGN method, we have used a pre-trained model, whereas for GraphPPIS we have trained the model ourselves.

We could not include in comparison the MaSIF program, due to 35 proteins of Dset_ 60 (over half) sharing similarity with the training dataset of MaSIF. It achieves an area under PR curve of 0.439, as reported in the GraphPPIS paper [52]. In spite of large similarity with its training data, the performance is comparable with that of Seq-InSite.

##### Comparison on Dset_ 315

The comparison on Dset_ 315 is presented in Table 6, with ROC and PR curves plotted in Figure 2(e)-(f). This is the only test where Seq-InSite comes in second place, behind GraphPPIS.

The MaSIF program, which, again, we could not include in comparison due to 104 proteins of Dset_ 315 (one third) sharing similarity with the training dataset of MaSIF, achieves an area under PR curve of 0.372, as reported in the GraphPPIS paper [52]. In spite of large similarity with their training data, their performance is lower than that of Seq-InSite.

Interestingly, when restricting the comparison to long proteins (length over 400), which are the most difficult to predict, Seq-InSite performs the best: area under the PR curve 0.211, compared to 0.184 for GraphPPIS and 0.145 for RGN.

##### Comparison on Dset_ 70

We use Dset_ 70 to include some comparison with several programs we could not run: AttentionCNN, DeepPPISP, EGRET, and HN-PPISP. All we can do is compare the computed area under the PR and ROC curves of Seq-InSite with those reported in the corresponding papers. We are unable to test GraphPPIS on all proteins in Dset_ 70 because GraphPPIS did not compute the feature set for all proteins in the combined dataset of Dset_ 72, Dset_ 164, and Dset_ 186.

The comparison on Dset_ 70 is presented in Table 7. Seq-InSite significantly outperforms the other methods, in spite of the fact that these are structure-based models.

**Table 7:**
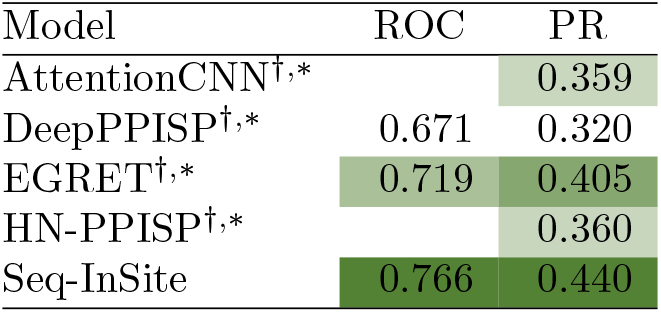
Performance comparison on Dset_ 70. The methods marked by ^*†*^ are structure-based. The methods marked by ^*^ were not trained by us; the data reported in their corresponding papers was used (see text for references). Darker colour indicates better performance.

| Model | ROC | PR |
| --- | --- | --- |
| AttentionCNN$\dagger,*$ | | 0.359 |
| DeepPPISP$\dagger,*$ | 0.671 | 0.320 |
| EGRET$\dagger,*$ | 0.719 | 0.405 |
| HN-PPISP$\dagger,*$ | | 0.360 |
| Seq-InSite | 0.766 | 0.440 |

### 3.4 Ablation study

We performed experiments aimed at clarifying the importance of various components and inputs of Seq-InSite. Combining the two architectural branches, MLP and LSTM, with the two input embeddings, MSA-transformer and ProtT5 XL (denoted MSA and T5 in these experiments), in all meaningful ways results in seven models, including the final ensemble. The results of testing all sub-models on Dset_ 448 are shown in Table 8, where the names of the models are self explanatory. We conclude from the results that the ProtT5 XL embeddings bring more information than the MSA-transformer ones. With fixed architecture, ProtT5 XL produces better performance. For the two architectures, MLP and LSTM, the comparison is less clear. LSTM is better when a single embedding is used, but MLP take the lead when both embeddings are used. Combining all four components produces the best results, and this is the Seq-InSite model.

**Table 8:** Ablation study using Dset_ 448.

| Model | Sens | Spec | Prec | Acc | F1 | MCC | ROC | PR |
| --- | --- | --- | --- | --- | --- | --- | --- | --- |
| MLP_MSA | 0.38 | 0.90 | 0.38 | 0.83 | 0.38 | 0.28 | 0.76 | 0.36 |
| MLP_T5 | 0.51 | 0.92 | 0.51 | 0.87 | 0.51 | 0.44 | 0.85 | 0.52 |
| LSTM_MSA | 0.42 | 0.91 | 0.42 | 0.84 | 0.42 | 0.33 | 0.78 | 0.40 |
| LSTM_T5 | 0.52 | 0.92 | 0.52 | 0.87 | 0.52 | 0.44 | 0.85 | 0.52 |
| MLP_MSA_T5 | 0.53 | 0.93 | 0.53 | 0.87 | 0.53 | 0.45 | 0.85 | 0.54 |
| LSTM_MSA_T5 | 0.51 | 0.92 | 0.51 | 0.87 | 0.51 | 0.44 | 0.85 | 0.53 |
| ENSEMBLE | 0.54 | 0.93 | 0.54 | 0.87 | 0.54 | 0.46 | 0.86 | 0.55 |

## 4 Evolutionary conservation

As an application of Seq-InSite’s predictive power, we analysed four different proteins. These are the alpha-subunit of the haemoglobin protein, the bacterial phosphoenolpyruvate carboxylase (PPC), the PPC’s four homologues in plants, and the human BRCA1 protein.

For the haemoglobin proteins, we used the same dataset as in DELPHI [25] to be able to compare the two results. The human haemoglobin was used for the query sequence (142 amino acids in length). For the other proteins, BLASTP was used to search for homologues. In the case of the bacterial PPC, the *E. coli* PPC protein sequence was used as the query (883 amino acids in length). In the case of the plant PPCs, the four *Arabidopsis thaliana* duplicate sequences were used as the query (967, 963, 968, 1032 amino acids in length). There are many isoforms of the human BRCA1 and so the canonical form (UniProtKB database entry P38398; 1863 amino acids in length) was used as the query. The search was restricted to the RefSeq protein database to ensure good quality hits. The top 100 hits were taken but excluded hits from Escherchia and Shigella in the case of PPC and excluded hits from the genus Homo in the case of BRCA1. For each set, an additional cluster BLASTP was done [45]. This database clusters together sequences within 90% identity and 90% length to other members of the cluster. This analysis of clustered sequences provides a taxonomically broader range of hits. Again the top 100 hits were taken. The query sequence, some selected homologues, and the two sets of sequences from the top 100 hits were combined. Sequences were aligned using MUSCLE [13]. Sequences that were identical, that contained unusually long stretches of inserted or deleted residues, or that were too distantly related to the query were manually trimmed with the aid of phylogenies constructed with IQ-TREE [34]. The alignment lengths for the proteins were: haemoglobin protein alignment length 308, bacterial phosphoenolpyruvate carboxylase alignment length 980, the PPC’s four homologues in plants alignment length 1282, 1254, 1029, 1341, and the BRCA1 protein alignment length 2157.

The results of Seq-InSite for the hemoglobin sequences are shown in Figure 3(a). The locations of important interfaces for the polypeptide interfaces to form a tetramer (db xref=”CDD:271278”) are shown with coloured arrows.

**Figure 3:**
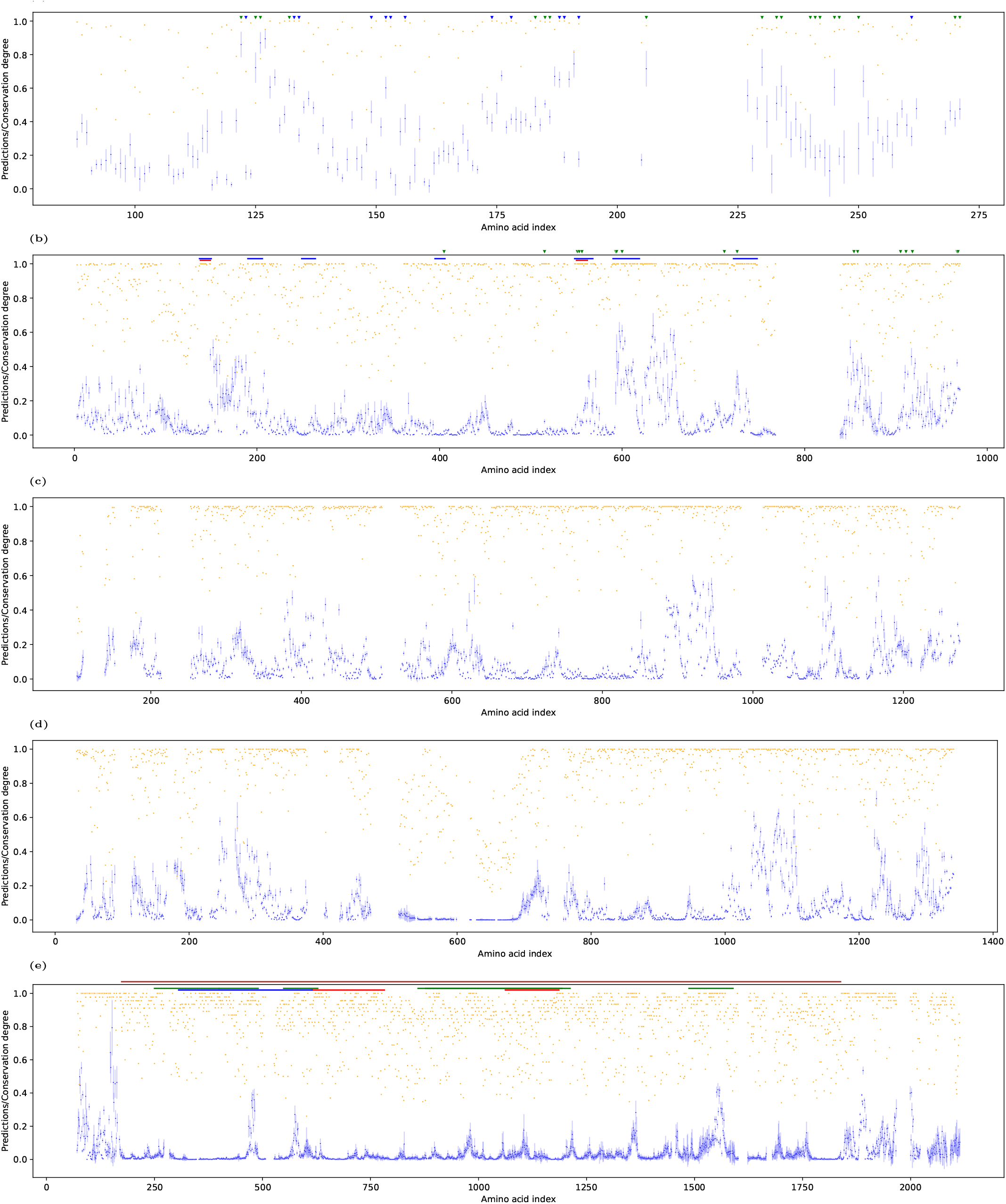
Interaction site predicted by Seq-InSite versus the degree in conservation. Seq-InSite predictions are shown in blue (mean in dark blue and variance in light blue) and degree of conservation in orange. The proteins are: (a) alpha haemoglobin – interfaces in green, binding sites in blue, (b) bacterial phosphoenolpyruvate carboxylase (PPC) – active sites in red, pepcarbxylase domains in blue and ligand binding sites in green, (c)-(d) PPC’s plant homologous, and (e) human BRCA1 protein – interaction colours: intrinsic disorder in brown, DNA binding in red, p53 binding in blue, protein binding in green.

It can be easily observed that the predicted PPI sites are in reasonable agreement with the actual sites involved in protein to protein interaction or protein sites binding to chemical moieties. The agreement is not perfect but primarily due to false negatives rather than false positives.

The bacterial PPC has a known three dimensional structure. The protein has two active sites, seven pepcarbxlase domains, and 17 ligand binding sites, highlighted in Figure 3(b) with coloured bars and arrows. The degree of agreement between these important sites and interaction probability within the plot can be observed. At the N-terminus there are none of these sites despite potentially high interaction probability which could be due to intra-peptide bindings.

Three of the plant PPCs (duplicates 1, 2 and 3) are very similar to the bacterial form. While there is no known three dimensional structure for plant PPCs, given their similarity it is doubtful that the structure would differ by very much and indeed, the predicted AlphaFold structure is similar. Since a BLASTP search using PPC1 query returned hits to PPC2 and PPC3, only the results for one (PPC1) is shown in Figure 3(c). The results for PPC4 are somewhat different and shown in Figure 3(d). A dotplot of PPC1 versus PPC4 shows a large insertion in PPC4 of approximately 100aa between amino acids 320 and 440 (466 and 718 in alignment) and shows poor similarity between the two proteins for the first 150 amino acids (235 aa in alignment). A PPI plot for PPC4 is shown in Figure 3(d) and suggests that, from this data, the large insertion is unlikely to be involved in protein interactions.

The human BRCA1 protein (UniProtKB P38398) is a difficult test for Seq-InSite. This protein is 83% in an intrinsically disordered state. As a result, AlphaFold can only make poorly supported predictions for the majority of the length of the protein. Seq-InSite, on the other hand, makes strong predictions at the amino terminus of the protein and less strongly supported predictions scattered throughout the length of the protein (Figure 3(e)). Like many other proteins with intrinsically disordered regions, the BRCA1 protein binds very promiscuously to a variety of other proteins. A few of its interactions are indicated in Figure 3(e) with coloured bars.

## 5 Discussion and conclusion

We have introduced a new program for protein interaction site prediction, Seq-InSite. In spite of the fact that Seq-InSite is sequence-based, it succeeded in matching or outperforming the much more accurate structure-based programs. In all our tests, Seq-InSite came in the first place, with a single exception, when it comes in second place. Despite this accuracy, the predicted sites (as exemplified in Figure 3) can not always be validated from the biological sequences alone. This does not mean that these predictions are wrong but rather that much still might be learnt from the sequences. Given the fact that sequences are much more readily available than structures, Seq-InSite can effectively replace structure-based methods for interaction site prediction.

It is interesting to note that the improved structure prediction from protein sequences, such as done by AlphaFold, may not bring the expected increase in use of structure in predicting various protein properties. This is due to the fact that the same mechanisms that enable the successful extraction and use of information from sequences to improve structure prediction can also help improve the prediction of other properties as well, directly from the sequence. Any information extracted from the sequence about structure and used subsequently as structure-based prediction, should be, at least in principle, used directly, bypassing structure prediction.

## 6 Availability

In order to make Seq-InSite widely available, we have built a web server at seq-insite.csd.uwo.ca, where the user provides protein sequences and receives the results by e-mail. The source code, including running instructions, pre-trained models, and all datasets used for training and testing, are available at github.com/lucian-ilie/seq-insite.

## 7 Author contributions statement

S.H. contributed to the methodology and analysis, designed the architectures, computed the training datasets, wrote the software, performed all tests, including installing and running competing methods, and wrote a draft of the manuscript.

G.B.G. designed and analyzed the experiments for the evolutionary conservation and wrote that section. L.I. proposed the problem, the ensemble architecture, the combined use of embeddings, analyzed the results, and wrote the final version of the manuscript.

## 8 Acknowledgments

Computations were performed on Compute Canada servers.

